# Spatio-dynamical decomposition: Joint spatial and dynamical criteria for FMRI Blind source separation

**DOI:** 10.64898/2026.09.25.754426

**Authors:** Sir-Lord Wiafe, Najme Soleimani, Tülay Adali, Vince D. Calhoun

## Abstract

Blind source separation (BSS) recovers latent sources from observed mixtures. In brain imaging, sources occur in two domains, spatial and temporal, so a criterion applied in one leaves the other unconstrained. Independent component analysis, the most used BSS tool, exploits diversity such as non-Gaussianity and sample dependence, but in one domain at a time. Methods that use both domains apply the same criterion to each and weight them against each other. We propose spatio-dynamical decomposition (SDD), scoring maps by non-Gaussianity and timecourses by dynamical independence: how much each source’s dynamics depend on the others. We normalize each score by how much it can improve on the data, eliminating the need for a weight. Across simulations and 130 naturalistic fMRI runs, SDD recovers sources distinguished by either maps or dynamics and reproduces both across disjoint episodes, while baselines trade off the two; map and timescale together also better predict stimulus tracking, with the largest gain for SDD. Together, these results show that separation is strongest when spatial and temporal domains each carry their own criterion.

## 1. INTRODUCTION

Blind source separation is standard wherever the measured channels are superpositions: speakers in microphone arrays, interference in communications, artifacts and networks in brain imaging [1, 2]. This matters most in brain imaging, where a measurement rarely isolates a single process [3]; thus, findings about cognition or disorder may reflect the mixing rather than the brain [4]. In functional MRI, these decompositions yield components, each with a spatial map and a timecourse, and both are interpreted: the map is matched to anatomy, the timecourse to task and behavior [5]. After a low-rank factorization, a BSS method chooses an invertible matrix using a criterion in the spatial domain on the maps, in the temporal domain on the timecourses, or in both.

Independent component analysis (ICA) is the most widely used of these estimators because, under its model, the sources are identifiable up to scale and permutation [6]. That identifiability can come from several diversity objectives such as non-Gaussianity, sample dependence, or both, and either form serves in either the spatial or the temporal domain [6]. For instance, spatial ICA might estimate the maps by maximizing their non-Gaussianity [5, 7], while weights-adjusted second-order blind identification (WASOBI) takes sample dependence into account and can be applied in either domain [8, 9]. Entropy-rate bound minimization (ERBM) scores them through entropy rate, which uses non-Gaussianity and sample dependence together; where as with any regular ICA approach both statistical property are applied to one domain at a time [10, 11]. Decompositions that score both domains at once exist, such as spatio-temporal ICA (stICA), which maximizes a weighted sum of spatial and temporal independence with a mixing parameter set by the user [12].

Across these, the selected criteria are applied within a single domain, whether one criterion or several, or a single criterion is applied across both. Which criterion suits a domain is a modelling choice, and the two domains are interpreted differently. A map is interpreted as a pattern over parcels, while a timecourse is interpreted as a process unfolding in time. We therefore propose spatio-dynamical decomposition (SDD), which uses a different form of diversity in each domain, each normalized by its attainable range on the data, eliminating a trade-off weight parameter. Here, maps are scored by the Infomax contrast, which favors sparse, localized patterns for spatial ICA to recover [5, 7]. Timecourses are scored by dynamical independence [13], measured by how much each component’s dynamics depend on the others. Here, we model those dynamics with a vector autoregression (VAR) over all components jointly, which describes macroscale brain activity better than nonlinear models [14]. We use second order, the lowest admitting complex-conjugate poles, thus the lowest order to represent a fluctuation decaying while it oscillates [15]. The same fit yields each component’s timescale [16].

We evaluate SDD in simulations and on naturalistic fMRI. In simulations, the sources are split into halves identifiable only from their maps or only from their dynamics. Our results show that methods restricted to one domain recover their own half and fail on the other; a joint method applying one criterion to both fails on the dynamic half, and SDD recovers both. Across 130 naturalistic fMRI runs from five subjects, baselines trade off reproducibility across domains, while SDD reproduces in both. Finally, a component’s map and dynamics together predict how closely it tracks speech better than either alone, and that gain is largest for SDD.

## 2. Methods

### 2.1. Model and identifiability

Let ***X*** ∈ ℝ^*V×T*^ hold one subject’s parcel time series, centered per parcel, with *V* parcels and *T* time points. A rank-*k* singular value decomposition gives an anchor (***S***_0_, ***A***_0_), where ***S***_0_ ∈ ℝ^*V*×*k*^ holds the *k* maps and ***A***_0_ ∈ ℝ^*k*×*T*^ their timecourses so that ***X*** ≈ ***S***_0_***A***_0_. For any invertible *k* × *k* matrix ***G***,

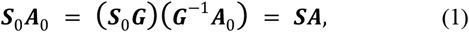

where ***S*** and ***A*** are the resulting maps and timecourses, respectively. The equality is exact, and it is the usual starting point for matrix decompositions. The anchor fixes the signal subspace but not the basis within it [1], so uniqueness follows only from constraints placed on ***S*** or on ***A***. Independence is one such constraint, and it can be imposed through non-Gaussianity, through sample dependence, or through both [6]. Estimating ***G*** is therefore the separation problem, and because the maps and the timecourses follow from that same ***G***, a criterion on either domain constrains one unknown rather than two (Fig. 1(a)). SDD constrains it from both domains, using different criteria in each.

**Fig. 1.**
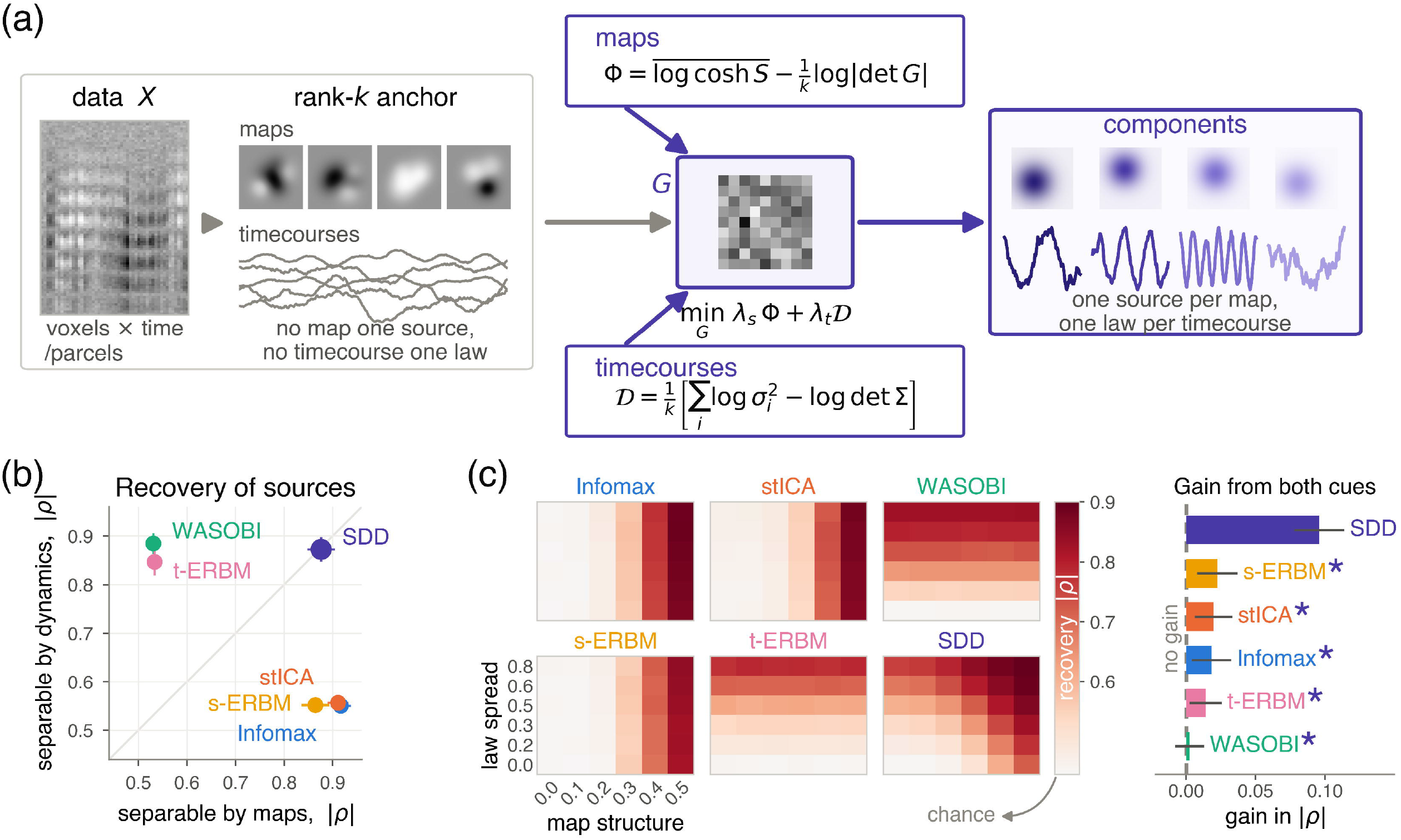
(a) The method: from a rank-k anchor, SDD estimates one k × k matrix G, read as maps and timecourses under spatial and dynamical criteria, each scaled by the range it can attain on the data. (b) Recovery of twenty sources, half separable only by their maps and half only by their dynamic laws; means over 60 replicates, bars 95% CIs. The diagonal, where only SDD sits, marks equal recovery of both halves. (c) Recovery over the plane of map structure and law spread, and the gain from using both domains at the strongest setting of each factor; SDD improves along both axes and gains most. * marks a baseline SDD beats (Wilcoxon signed-rank, FDR-corrected q < 3.2 × 10^17^).

### 2.2. Spatial criterion

The anchor is whitened, so its maps have unit variance and are uncorrelated, and the criterion measures shape rather than power. The resulting maps are ***S*** = ***S***_0_***G*** ∈ ℝ^*V×k*^, where ***S***_0_ denotes the whitened anchor and *S*_*vi*_ is the entry of ***S*** giving the loading of parcel *v* on component *i*. The maps are scored by

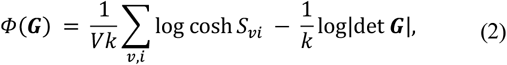

which is the Infomax cost [17] which is equivalent to the negative log-likelihood of a super-Gaussian density and the log-determinant of the transformation, divided by *k* so that both terms are per-component quantities.

### 2.3. Dynamical criterion

The timecourses are ***A*** = ***G***^−2^***A***_0_ ∈ ℝ^*k×T*^, whose column ***a***_*t*_ ∈ ℝ^*k*^ holds the *k* component values at time *t*. A second-order vector autoregressive model, VAR(2), is fitted to ***A***,

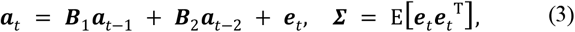

where Σ is the innovation covariance of the joint fit of all *k* components. Regressing each component on its own past alone gives the residual variance, 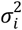. The gap between the two fits then scores the timecourses,

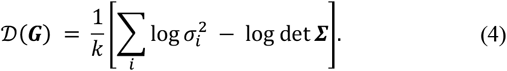

Since Σ = ***G***^−2^Σ_0_***G***^−*T*^, *D* equals 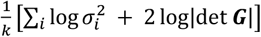 up to a constant, the Gaussian mutual-information rate among the components [18]. Adding predictors cannot increase the residual variance, so 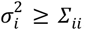, and log det Σ ≤ ∑_*i*_log Σ_*ii*_ by Hadamard’s inequality. Hence *D* ≥ 0, with equality when each component is predicted as well by its own past as by the whole system, with uncorrelated innovations [18], the dynamical independence condition [13]. In the taxonomy of diversity, this is a form of sample dependence [6], obtained here from a fitted dynamical model rather than a lagged statistic. Large values mean the dynamics are spread across the set, so predicting one component requires the others. For a given ***G***, the ***B***_1_, ***B***_2_ and Σ follow in closed form from the lag moments of ***A***, so each evaluation solves them directly, leaving ***G*** as the only unknown.

### 2.4. Objective and estimation

SDD estimates

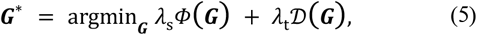

where λ_s_ and λ_t_ place the two criteria on a common scale. Each is the reciprocal of the range its criterion can attain on the data, The weights are

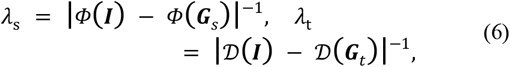

where ***I*** is the identity, and ***G***_*s*_ and ***G***_*t*_ minimize their respective criteria. A criterion that varies little across a dataset is up-weighted, while one that varies more is down-weighted, so neither dominates. SDD therefore pairs non-Gaussianity in the maps with dynamical independence in the timecourses, a different form of diversity in each domain [6]. With λ_=_ = 0, Eq. 5 reduces to the Infomax objective minimized over the same free ***G***, so the Infomax baseline is exactly the spatial-only ablation of SDD. The objective is minimized over the *k*^2^ entries of ***G*** by L-BFGS [19] from ***G*** = ***I***, using analytic gradients for both terms. Eight random orthogonal starts reach the same solution, with matched map |ρ| ≥ 0.97 and objectives within 0.3%. Once the lag moments are formed at *O*(*Tk*^2^), each evaluation costs *O*(*Vk*^2^ + *k*^3^) independent of *T*.

## 3. DATA AND EXPERIMENTS

### 3.1. Simulation

Twenty sources are mixed into V parcels over T time points, with V, T, and the noise level (NL) drawn independently per replicate over V = 400–1500, T = 200–1200, and NL = 0.02–0.40. Two factors determine source identifiability. Map structure controls focality, interpolating between spatially unstructured noise and sparse parcel supports drawn from a shared pool. Law spread controls how much the sources’ dynamical laws differ, interpolating between one law shared by all sources and a different law for each. Laws are impulse responses from five families: AR(2), AR(4), ARMA, fractional long-memory, and damped oscillator. SDD fits only VAR(2), so four families are mis-specified.

We report two designs. First, the factors are separated: half the sources are identifiable only from their maps and half only from their laws, forming map-separable and law-separable halves, with 60 replicates per condition. Second, both factors apply to every source and vary over six levels each, map structure from 0 to 0.5 and law spread from 0 to 0.8, giving 36 conditions with 40 replicates each. The zero condition is a design check: no source is identifiable, so all methods should perform at chance, which is the mean recovery over 20 random rotations. Recovery is the mean absolute correlation between each true source map and its Hungarian-matched estimate. At the strongest setting of each factor, we measure the gain from using both domains against the better single domain.

### 3.2. fMRI data

We use the Friends naturalistic movie-viewing dataset [20]: five subjects, thirteen episodes each recorded as two runs, for 26 runs per subject (130 runs in total), at TR = 1.49 s with about 470 volumes per run. Each run was parcellated with the Schaefer 1000-parcel atlas (V=1000) [21], nuisance-regressed using its own fMRIPrep confounds [22], including 24 motion parameters, five aCompCor components [23], and linear and quadratic trends, then standardized per parcel. First, reproducibility is tested across disjoint halves of each subject’s episodes, with both runs of an episode kept in the same half, using five different partitions. Each half is decomposed independently, and components are matched across halves by Hungarian assignment on the absolute correlation between their maps. Second, we test cognitive relevance using speech as an external criterion. Its timing and cortical locus are known, allowing us to test whether components track an interpretable stimulus. From the time-aligned transcripts, we construct a word-presence regressor that is one at each TR containing a word onset and zero otherwise, then convolve it with a canonical double-gamma hemodynamic response [24]. We define speech coupling as the absolute correlation between a component timecourse and this regressor. Components are estimated leave-one-run-out: each decomposition is fitted on 25 of a subject’s 26 runs and measured on the one held out. Auditory share is the fraction of parcels with |*z*| > 2 whose Schaefer labels are Temp, TempPole, TempOcc, TempOccPar, FrOperIns, or ParOper. Auditory share and timescale are used to predict speech coupling, with models fitted on four subjects and scored on the fifth.

### 3.3. Baselines and settings

We compare against spatial Infomax [17], which maximizes the output entropy of the maps, favoring super-Gaussian maps; stICA [12], which maximizes a weighted sum of spatial and temporal independence; WASOBI [8, 9], which jointly diagonalizes lagged covariances of the timecourses under asymptotically optimal weighting; and ERBM [10, 11], which minimizes entropy rate and is applied in both domains, to the maps (s-ERBM) and timecourses (t-ERBM). All methods estimate the same number of components from the same anchor. Reproducibility is assessed in both domains: directly for the maps and dynamically through each component’s timescale, measured by ACW-50, the lag at which autocorrelation falls to one-half [16]. ACW-50 is model-free and not optimized by any method here. In simulations, stICA’s mixing weight, WASOBI’s AR order, and ERBM’s filter length each take the value maximizing that method’s ground-truth recovery; SDD and Infomax have no tunable parameters. On real data, where no answer key exists, all use fixed settings: stICA at α = 0.5, ERBM at its default filter length of 11 [11], and WASOBI at AR order 11 to match. Unless stated otherwise, real-data analyses use *k*=40 components; reproducibility is also evaluated for *k* from 10 to 100.

## 4. RESULTS

### 4.1. Simulation

Fig. 1(b) reports the first design, where the methods behave as expected. Infomax, the spatial-only ablation of SDD, stICA, and s-ERBM recover the map-separable half at 0.92, 0.91, and 0.86, but fall to 0.55, 0.56, and 0.55 on the law-separable half. WASOBI and t-ERBM show the reverse, reaching 0.88 and 0.85 on the law-separable half and 0.53 on the other. SDD alone exceeds 0.87 on both, at 0.876 and 0.873, and alone sits on the diagonal. stICA fails on the law-separable half despite estimating both domains, reflecting the cost of reusing one criterion. Fig. 1(c) reports the second design, sweeping map structure and law spread together. When both are absent, no method exceeds chance (0.454), confirming that the design carries no incidental structure. Elsewhere, each baseline improves along one axis only, whereas SDD improves along both. At the strongest setting of each factor, the gain from using both domains is 0.096 for SDD against 0.002 to 0.022 for the baselines, exceeding every one of them (Wilcoxon signed-rank, FDR-corrected, *q* < 3.2 × 10^−7^).

### 4.2. Reproducibility on real data

Fig. 2(a) shows four SDD components at k = 40, recovered from disjoint halves of one subject’s episodes and chosen to span its timescale range. Their maps correlate across halves at |ρ| ≥ 0.92, timescales agree within 0.3 s, and autocorrelation functions correlate at |ρ| ≥ 0.80. Both descriptions therefore survive independent estimation from different episodes. Fig. 2(b, c) extends this analysis across subjects, partitions, and model sizes. For maps, SDD is highest throughout, reaching 0.802 at k = 40 and 0.612 at k = 100. It is matched only by Infomax and exceeds the dynamics methods by 0.08–0.18. For timescales, SDD is intermediate at small k but overtakes the baselines as the model grows, reaching 0.950 at k = 100 versus 0.942 for the best baseline, while stICA falls to 0.096. Fig. 2(d, e) gives the paired comparison. SDD minus each baseline is positive in 30 of 30 map cells and 25 of 30 timescale cells, with significant differences in 38 of 60 (subject as a random effect, FDR-corrected across all 60 tests). No baseline is competitive in both domains: the map-based methods lose on timescales, while the dynamics methods lose on maps.

**Fig. 2.**
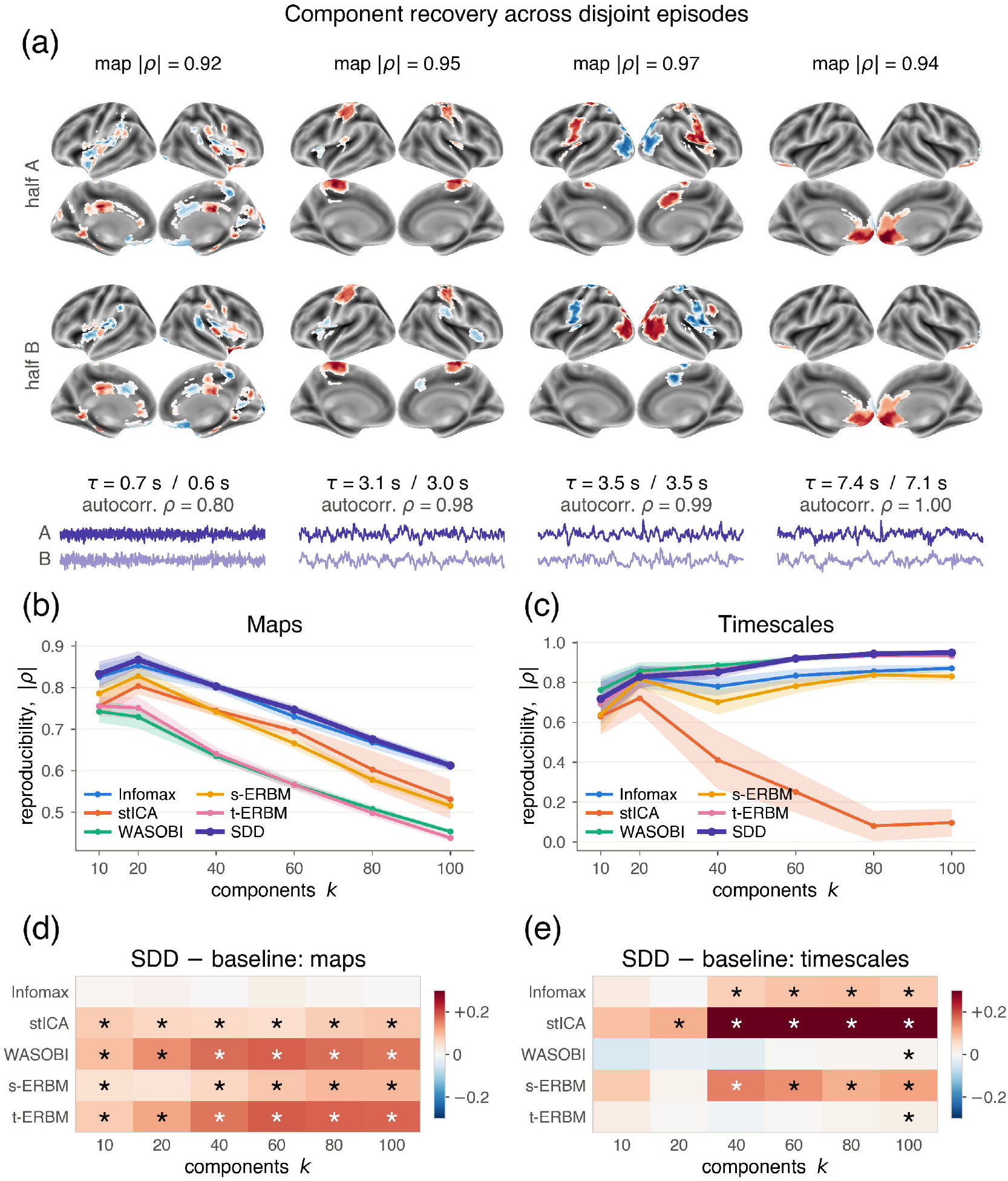
Split-half reproducibility on real data. (a) Four SDD components at k = 40 spanning its timescale range: maps from halves A and B, a timecourse from each, and their agreement (maps |ρ| ≥ 0.92, timescales within 0.3 s, autocorrelation |ρ| ≥ 0.80). (b, c) Map and timescale reproducibility against k; SDD is highest in maps and overtakes in timescales as k grows. (d, e) SDD minus each baseline per domain, averaged over the 25 subject × partition cells. * marks a cell surviving FDR correction over the 60 tests in the two panels, with subject as a random effect (q < 0.05).

### 4.3. Stimulus tracking by the decomposed components

A reproducible component need not be informative, so Fig. 3 asks whether reproducible components track speech. We use each component’s auditory share and timescale to predict its speech coupling, the strength of its correspondence with the speech regressor. Both together predict better than either alone for every method, and the gain is largest for SDD at 0.024 against 0 to 0.016 for the baselines, exceeding each of them (Fig. 3(a)). SDD’s gain is also the most consistent across subjects; its lowest subject exceeds the median subject of every baseline (hollow circles, Fig. 3(a)). Fig. 3(b) tests anatomical plausibility. Among the top decile by speech coupling, 29.7% of an SDD component’s parcels fall in the auditory cortex, versus 19.3–24.8% for the baselines. Across all components, the methods are indistinguishable at 15.0–16.5%, so the enrichment is specific to speech-coupled components. SDD therefore isolates the auditory response more cleanly than the baselines do.

**Fig. 3.**
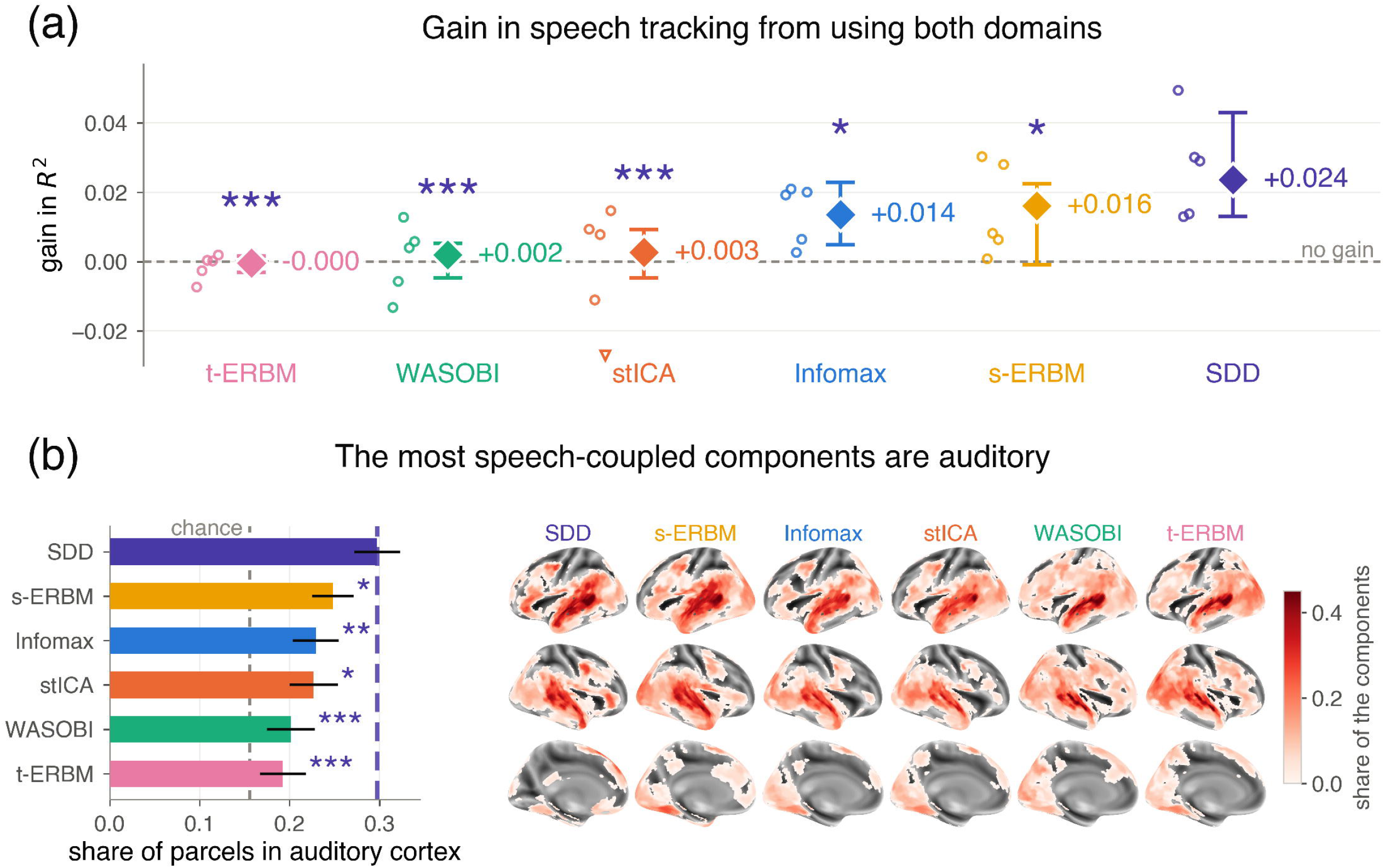
Speech tracking by the recovered components, each described by its share of parcels in the auditory cortex and its timescale. (a) Gain of both coordinates over the better single one, in leave-one-run-out prediction of speech coupling; diamonds are medians over 130 runs with bootstrap 95% intervals, circles are the five per-subject medians. The gain is 0.024 for SDD against 0 to 0.016 for the baselines. (b) The top decile of components by speech coupling: their share of parcels in the auditory cortex (SDD 29.7% against 19.3–24.8%, chance 15.6%) and their maps per method, on one scale. * marks SDD above a baseline, FDR-corrected (* q < 0.05, ** q < 0.01, *** q < 0.001).

## 5. CONCLUSION

We propose spatio-dynamical decomposition, which constrains the source separation transformation from both domains rather than one, with a different criterion in each. Each score is divided by the range it can attain on the data, so the two are comparable without a trade-off weight. In simulations, SDD recovers sources that only their maps or only their dynamics separate, while each baseline recovers one and fails on the other. On 130 naturalistic fMRI runs, its components reproduce across disjoint episodes in both domains, and their maps and timescales together predict speech tracking better than either alone, by more than any baseline. Identifiability of the combined objective is not established here; the support is empirical, across recovery, reproducibility, and stimulus tracking. In SDD, the criteria are replaceable, chosen to suit what each domain represents rather than fixed by the construction. SDD therefore offers a general framework for source separation in which each domain carries the criterion suited to it.

